# Leaf-level metabolic changes in response to drought affect daytime CO_2_ emission and isoprenoid synthesis pathways

**DOI:** 10.1101/2022.04.29.490001

**Authors:** S. Nemiah Ladd, L. Erik Daber, Ines Bamberger, Angelika Kübert, Jürgen Kreuzwieser, Gemma Purser, Johannes Ingrisch, Jason Deleeuw, Joost van Haren, Laura K. Meredith, Christiane Werner

**Affiliations:** Ecosystem Physiology, University of Freiburg, Georges-Köhler-Allee 053/054, 79110 Freiburg, Germany; Department of Environmental Sciences, University of Basel, Bernoullistrasse 30, 4056 Basel, Switzerland; School of Chemistry, The University of Edinburgh, Joseph Black Building, David Brewster Road, Edinburgh, EH9 3FJ, UK; UK Centre for Ecology & Hydrology, Bush Estate, Penicuik, EH26 0QB, U K; Department of Ecology, University of Innsbruck, Sternwartestrasse 15, 6020 Innsbruck, Austria; Biosphere 2, University of Arizona, 32540 S. Biosphere Rd., Oracle, AZ, 85739, USA; Honors College, University of Arizona, 1101 E. Mabel Street, Tucson, AZ, 85719, USA; School of Natural Resources and the Environment, University of Arizona, Tucson, AZ, USA

**Keywords:** daytime respiration, drought, GC-IRMS, isoprenoids, legumes, position-specific, isotope labeling, PTR-ToF-MS, tropical plants, volatile organic compounds

## Abstract

In the near future, climate change will cause enhanced frequency and/or severity in terrestrial ecosystems, including tropical forests. Drought responses by tropical trees may affect their carbon use, including production of volatile organic compounds (VOCs), with unpredictable implications for carbon cycling and atmospheric chemistry. It remains unclear how metabolic adjustments by mature tropical trees in response to drought will affect their carbon fluxes associated with daytime CO_2_ production and VOC emission. To address this gap, we used position-specific ^13^C-pyruvate labeling to investigate leaf CO_2_ and VOC fluxes from four tropical species before and during a controlled drought in the enclosed rainforest of Biosphere 2. Overall, plants that were more sensitive to drought had greater reductions in daytime CO_2_ production. Although daytime CO_2_ production was always dominated by non-mitochondrial processes, the relative contribution of CO_2_ from the tricarboxylic acid cycle tended to increase under drought. A notable exception was the legume tree *Clitoria fairchildiana*, which had less anabolic CO_2_ production than the other species even under pre-drought conditions, perhaps due to more efficient refixation of CO_2_ and anaplerotic use for amino acid synthesis. *C. fairchildiana* was also the only species to allocate detectable amounts of ^13^C label to VOCs, and was a major source of VOCs in the Biosphere 2 forest. In *C. fairchildiana* leaves, our data indicate that intermediates from the mevalonic acid pathway are used to produce the volatile monoterpene trans-β-ocimene, but not isoprene. This apparent crosstalk between the mevalonic acid and methylerythritol phosphate pathways for monoterpene synthesis declined with drought. Finally, although trans-β-ocimene emissions increased under drought, it was increasingly sourced from stored intermediates and not *de novo* synthesis. Unique metabolic responses of legumes may play a disproportionate role in the overall changes in daytime CO_2_ and VOC fluxes in tropical forests experiencing drought.

## Introduction

Due to anthropogenic climate change, the frequency and/or intensity of droughts are expected to increase, particularly in tropical regions (Douville et al., 2021). The responses of tropical forests to droughts can have important feedbacks on the global carbon cycle, as these forests represent a significant carbon sink today, but may lose that capacity as warming and droughts increase (Mitchard 2018; Hubau et al., 2020). Additionally, tropical forests are an important source of volatile organic compounds (VOCs) to the atmosphere (Guenther et al., 2006). The overall fluxes and molecular composition of VOCs emitted from tropical forests are also likely to change with increased drought (Peñuelas and Llusia, 2003), with further implications for atmospheric chemistry (Atkinson and Arey, 2003; Claeys et al., 2004; Lelieveld et al., 2008). It is therefore important to better characterize and understand changes in carbon fluxes, including allocation to VOCs, on an ecosystem and individual tree scale in tropical forests in response to drought.

In depth investigations of drought in natural forests remain challenging, since the timing of severe droughts is difficult to predict in advance and it is rarely practical to leave extensive analytical equipment in place for long periods of time in the hopes of capturing an extreme event. Model ecosystems such as the enclosed tropical rainforest in the Biosphere 2 (B2) experimental facility in Oracle, AZ, USA, thus provide a valuable means to experimentally investigate drought impacts in a near-natural setting (Rascher et al., 2004; Pegoraro et al., 2006; Evaristo et al., 2019; Werner et al., 2021). In 2019, the Biosphere 2 Water Atmosphere and Life Dynamics (B2WALD) campaign made use of this unique experimental ecosystem to investigate drought from molecular to ecosystem scales by subjecting it to a two-month drought accompanied by intensive *in situ* measurements and sampling of different ecosystem components before, during, and after drought (Werner et al., 2021). Among other findings, initial analyses from the B2WALD campaign demonstrated that overall ecosystem fluxes of water, CO_2_, and VOCs were influenced in different ways by four distinct functional plant groups: drought-sensitive and drought-tolerant canopy trees, and drought-sensitive and drought-tolerant understory plants (Werner et al., 2021, for further details, see **Figure 1**). While it is clear that overall changes in ecosystem CO_2_, water, and VOC fluxes were driven by differing responses from these four plant functional groups during the B2WALD drought, the specific metabolic adjustments within each of these four plant groups have not yet been investigated. In particular, changes in leaf gas exchange, water use efficiency, and partitioning of carbon between catabolic and anabolic processes, including the biosynthesis of VOCs, could potentially provide valuable insight into how and why plant functional groups adjusted their metabolism towards drought.

**Figure 1:**
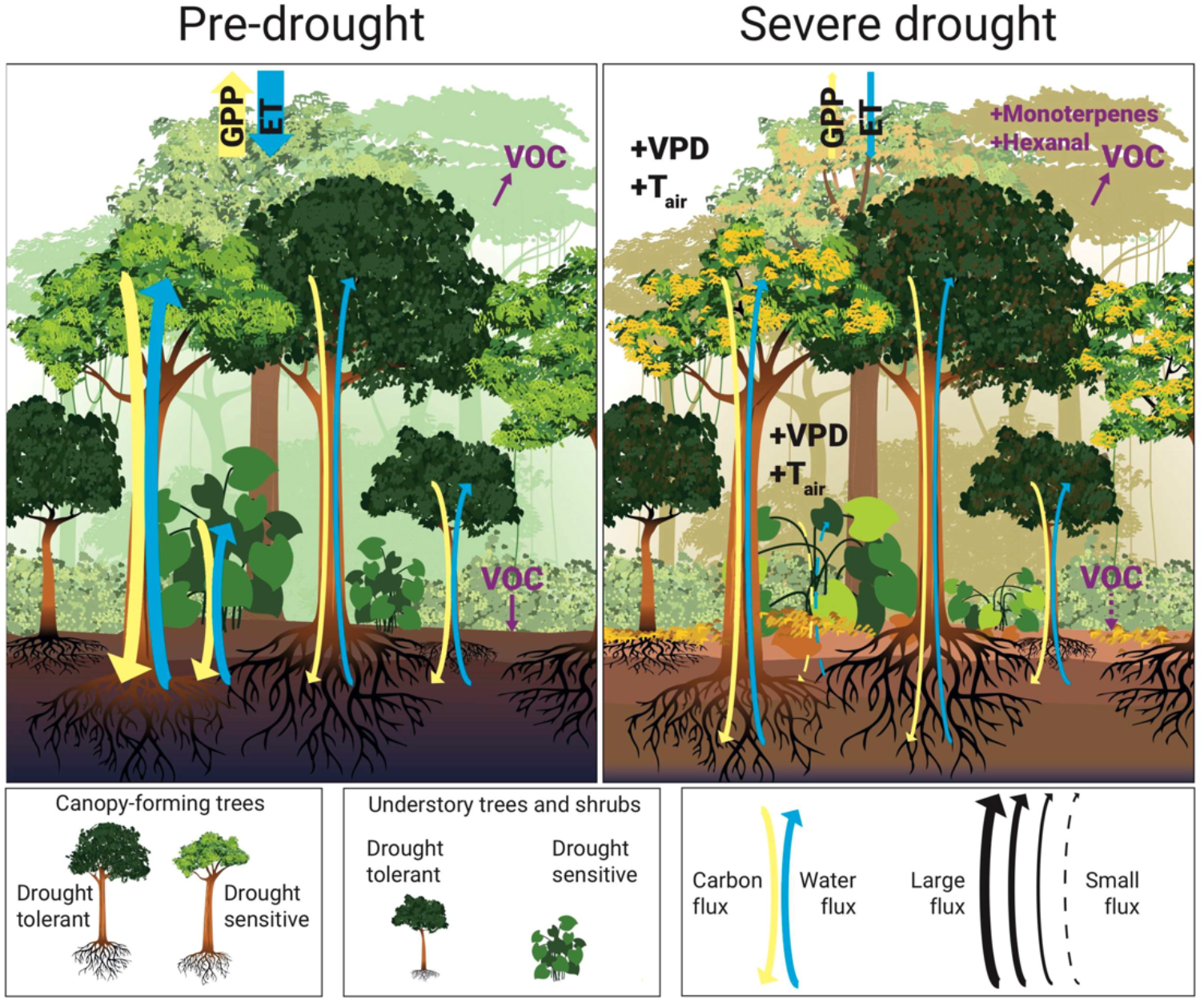
Schematic representation of drought-responses by four plant functional groups identified in the B2 WALD drought, modified from Werner et al., 2021. Drought-sensitive canopy trees were responsible for the majority of ecosystem fluxes before the drought, but rapidly dropped leaves and reduced transpiration demand as upper soil layers dried. Their drought-tolerant counterparts never moved water through their stems as quickly, but had much smaller reductions in activity and leaf water potential in response to drought. Meanwhile, in the understory, drought tolerance was largely driven by microclimate variability, with shaded plants having low fluxes of carbon and water throughout the study period that did not change with drought and were most likely determined by limited light availability. Understory plants in locations with more light availability, either from canopy gaps or windows, were much more sensitive to drought and were the most stressed plants by the end of the experimental drought.

To investigate metabolic adjustments towards drought and how they relate to the diverse drought responses of different plant functional groups, we measured the incorporation of ^13^C from position-specific pyruvate labels into CO_2_ and VOCs before and during the B2WALD drought. This approach is based on the fact that different carbon atoms within the central metabolite pyruvate are used for distinct biochemical processes. For example, carbon atoms from the C2 and C3 positions of pyruvate are converted to CO_2_ by respiration in the tricarboxylic acid (TCA) cycle (**Figure 2, step 1**) (Tcherkez et al., 2005; Priault et al., 2009), while carbon atoms from the C1 position of pyruvate are also released due to TCA cycle activity (**Figure 2**) (Tcherkez et al., 2008; Priault et al., 2009). However, numerous non-mitochondrial anabolic processes also decarboxylate the C1 position of pyruvate, while the C2 and C3 position are incorporated into biosynthetic products (**Figure 2**) (Graus et al., 2004; Schnitzler et al., 2004; Jardine et al., 2010). Together, these processes cause greater ^13^C enrichment of CO_2_ following labeling with ^13^C1-pyruvate than ^13^C enrichment of CO_2_ following labeling with ^13^C2-pyruvate during the day, when the TCA cycle is downregulated (Tcherkez et al., 2008; Priault et al., 2009). The relative difference in ^13^C enrichment of CO_2_ following labeling with each type of pyruvate is thus broadly indicative of the relative significance of CO_2_ production from non-mitochondrial anabolic processes (Priault et al., 2009; Werner et al., 2009; Tcherkez et al., 2012; Fasbender et al., 2018).

**Figure 2:**
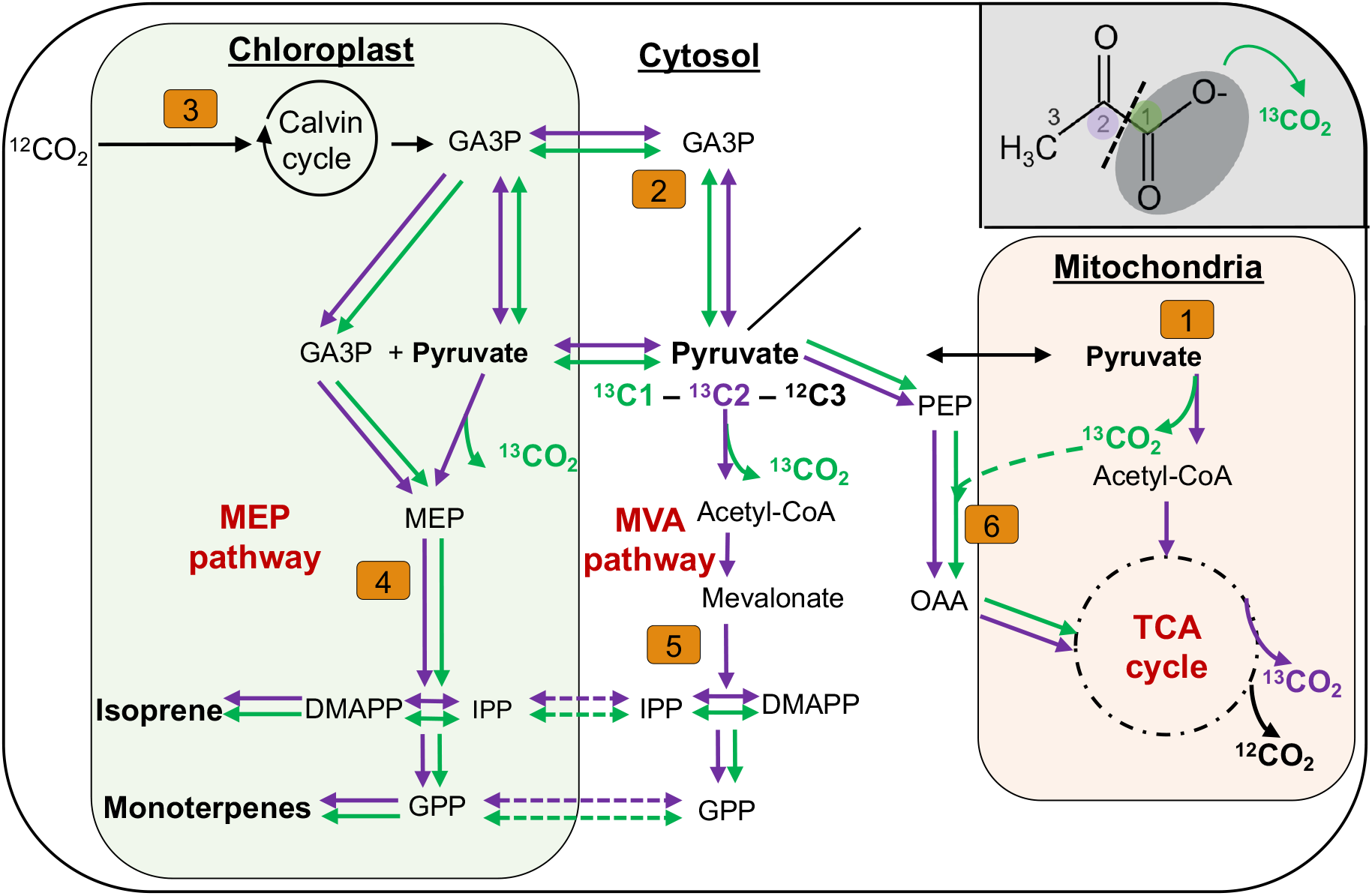
Schematic representation of important processes for the synthesis of isoprenoids and production of CO_2_ within plant cells, with several intermediate steps removed for clarity. Green arrows represent possible movement of carbon from the C1 position of pyruvate and purple arrows represent movement of carbon from the C2 position of pyruvate. Numbers represent the relevant steps discussed. 1: Transport of pyruvate into mitochondria, followed by initial decarboxylation of the C1-position and entrance into the TCA-cycle. 2: Conversion of pyruvate to GA-3-P and transport into the chloroplast. Pyruvate might also be transported directly to the chloroplast. All carbon positions of pyruvate remain in both cases. 3: Entrance of CO_2_ into the Calvin cycle and addition of GA-3-P and other assimilation products into the metabolic pool of the chloroplast. 4: Initial reaction of pyruvate and GA-3-P to form DMAPP and IPP via MEP pathway, followed by reaction of two DMAPP/IPP molecules to produce GPP. As GA-3-P is not decarboxylated in this reaction, all carbon positions of the converted GA-3-P remain in the produced isoprene and monoterpenes. Pyruvate on the other hand is decarboxylated, resulting in only the C2-position remaining in these products 5: Entrance of pyruvate into MVA pathway and initial decarboxylation of the C1-position of pyruvate to form DMAPP, IPP and subsequently GPP in the cytosol. After transport of these metabolites incorporating the ^13^C2-label into the chloroplast, DMAPP and IPP can either enter isoprene or monoterpene synthesis, while GPP is used to synthesize monoterpenes. 6: Anaplerotic replenishment of OAA to the TCA cycle from PEP and CO_2_, which can support the production of carbon backbones for N-containing compounds. Abbreviations: TCA, tricarboxlic acid; GA-3-P, glyceraldehyde 3-phosphate; DMAPP Dimethylallyl pyrophosphate; IPP, isopentenyl pyrophosphate; MEP, methylerythritol phosphate; GPP, geranyl pyrophosphate; MVA, mevalonic acid; PEP, phospho*enol*pyruvate; OAA, oxaloacetate.

Position-specific ^13^C labeling can also be employed to study the synthesis of various plant metabolites, including VOCs (Schwender et al., 1996; Lichtenthaler et al., 1997; Schuhr et al., 2003; Fasbender et al., 2018; Werner et al., 2020). This approach can be particularly helpful for understanding the biosynthesis of both volatile and non-volatile isoprenoids, and for assessing the role of metabolic crosstalk between the cytosolic mevalonic acid (MVA) and the plastidic methylerythritol phosphate (MEP) pathways for isoprenoid biosynthesis (**Figure 2, step 4 & 5**) (Schuhr et al., 2003; Massé et al., 2004; Werner et al., 2020; Ladd et al., 2021).

While position-specific ^13^C labeling is well established in laboratory experiments on potted plants, no study has used this approach to assess metabolic adjustments of mature plants in response to drought in a natural ecosystem. Based on the initial characterization of four functional groups in the B2 rainforest, we labeled leaves from *Clitoria fairchildiana* R.A. Howard (drought-sensitive canopy)*, Phytolacca dioica* L. (drought-tolerant canopy), *Hibiscus rosa sinensis* L. (drought-tolerant understory), and *Piper auritum* Kunth (drought-sensitive understory) with position-specific ^13^C-pyruvate before and during the B2WALD drought to address the following hypotheses: (1) drought-sensitive plants with large reductions in assimilation and transpiration also produce and release less CO_2_ from pyruvate, as their overall metabolism slows, (2) the majority of cytosolic pyruvate is used for non-mitochondrial processes in the light under control conditions, but the difference with use in the TCA cycle declines as drought responses intensify, and (3) for isoprenoid emitting plants, the proportion of isoprenoids synthesized from cytosolic pyruvate increases under drought in drought-sensitive plants as the supply of fresh photosynthate available for biosynthesis declines. Together, this approach allows us to evaluate how allocation of pyruvate changed under drought, and how these metabolic adjustments relate to the diverse integrated drought responses of each of these four tropical plant species.

## Materials and Methods

### Drought campaign

This study took place within the context of the broader B2WALD campaign, described in detail by Werner et al. (2021). The B2WALD campaign made use of the enclosed rainforest that has been growing continuously for over 30 years at the B2 research facility, located north of Tucson, AZ, USA. The enclosed system is equipped with sprinklers that control precipitation within the forest, thus enabling full control over the start and end dates for the drought experiment. Additionally, temperature and ambient CO_2_ concentrations within the forest are elevated compared to natural tropical rain forests, making this system a potential analogue for future climate scenarios (Lloyd and Farquhar, 2008). In 2019, this mature ecosystem was subjected to a 65-day drought, with intensive monitoring of gas fluxes from leaves, stems, soils, and roots throughout a 110-day period before, during, and after drought (Werner et al., 2021). Under non-drought conditions, the Biosphere 2 rainforest receives 1600 mm/year, distributed by overhead sprinklers three times per week. Following the final pre-drought rain event on the night of 7 October 2019, we turned off the sprinkler system and did not add additional water to the system until 2 December 2019. Daytime temperature within the canopy of the Biosphere 2 rainforest ranged from 23.3 to 33.5 °C during the pre-drought period and 22.2 to 34.8 °C during the drought. Corresponding daytime temperature ranges for the understory were 21.3 - 26.6 °C and 22.2 - 29.6 °C, respectively. maximum photosynthetically active radiation (PAR) values outside the enclosure declined from 1685 to 1016 μmol m^-2^ s^-1^ (Werner et al., 2021).

During the first four weeks of the B2WALD drought, surface soil moisture was almost completely depleted, and the atmospheric vapor pressure deficit increased in both the canopy and the understory, but at a higher rate in the canopy (Werner et al., 2021). In normal years there is no seasonal variability in precipitation within the Biosphere 2 rainforest, and the ecosystem-scale changes in water and carbon fluxes were much stronger than seasonal changes associated with reduced external light availability. Changes in water fluxes by individual plants were also clearly driven primarily by drought and not light availability, as the trees’ sap flow increased immediately when rain first returned to the system in mid-December, when ambient light levels were at their annual minimum.

### Gas sampling system and analyzers

For all target species (*Clitoria fairchildiana, Piper auritum, Phytolocca dioica* and *Hibiscus rosa sinensis*), we enclosed whole leaves in species-specific, flow-through ventilated cuvettes made from FEP film and PFA tubing as described in Kübert et al. (2023). One end of the leaf cuvette was sealed around the petiole, while the other end was sealed around the incoming and outcoming PFA tubes. For *H. rosa sinensis*, we used cuvettes that enclosed several leaves on terminal branches. This was necessary because it was difficult to avoid embolism when cutting the petiole of single *H. rosa sinensis* leaves. We equipped each cuvette with a data logger and a light sensor that recorded light levels at an angle parallel to the leaf’s surface, as well as a thermocouple that recorded leaf surface temperature. Leaf temperature inside the leaf cuvettes did not exceed daytime ecosystem temperature maxima reported in Werner et al. (2021) (**Table S1**).

We continuously supplied each cuvette with a mix of VOC- and CO_2_-free air produced by a zero air generator (Aadco Instruments, Inc., Cleves, OH, USA) and a constant amount of CO_2_ (~500 ppm, δ^13^C ~ −10 ‰) from a gas tank. This relatively high CO_2_ concentration was selected to match the average ambient concentrations within the Biosphere 2 rainforest. Each cuvette was equipped with a fan to guarantee efficient air mixing. The outflow from the cuvette was connected to a Teflon T-piece and ball valve that allowed us to sample a portion of the flow onto cartridges for offline VOC analyses (details below). Additional flow was diverted to three analyzers: (1) a proton transfer reaction – time of flight mass spectrometer (PTR-TOF-MS 4000ultra, Ionicon Analytic, Innsbruck, Austria) for VOC fluxes, (2) a carbon isotope laser spectrometer (Delta Ray IRIS, Thermo Fisher Scientific, Bremen, Germany) for concentrations and stable isotopes of CO_2_, and (3) a water isotope cavity ring-down spectrometer (L2120i water isotope analyzer, Picarro Inc., Santa Clara, CA, USA) for concentrations and stable isotopes of water vapor. Details of the gas sampling system, analyzers, and the calibration of different analyzers used during the B2WALD campaign are provided in Werner et al. (2021).

### Position-specific ^13^C-pyruvate labeling

We conducted position-specific ^13^C-pyruvate labeling during two time periods: pre-drought (12 – 28 September 2019) and drought (4 – 16 November, 2019; after 27 – 39 days without rain), using a protocol similar to that of Fasbender et al. (2018). For canopy trees, we did the labeling during the morning hours, when overall photosynthesis tends to be highest in the B2 rainforest, as hot temperature in the sunlit canopy induce early stomatal closures (Rascher et al., 2004; Rosolem et al., 2010). For the cooler understory, we primarily labeled in the afternoon. Leaves were installed in cuvettes at least one day prior to the labeling event. Immediately prior to labeling, we monitored the cuvette for 15 minutes to ensure that leaf fluxes of CO_2_ and VOCs were stable. We then cut the leaf petiole and rapidly placed it under water, and cut it again to avoid embolism. The petiole remained in a 1.5 mL centrifuge tube filled with water for five more minutes so that the fluxes could stabilize after cutting. Subsequently, we exchanged the centrifuge tube with one containing a 10 mM solution of either ^13^C1- or ^13^C2-pyruvate (99 atom % ^13^C, Cambridge Isotope Laboratories, Andover, MA, USA). We used the mass decline of the centrifuge tube over time to calculate the label uptake. In most cases, leaf fluxes of H2O, CO_2_, and VOCs returned to the levels they had while attached to the plant soon after cutting. The few leaves that did not stabilize at or near their previous gas fluxes after being placed in the label solution were excluded from subsequent analyses.

We collected the leaf after each measurement to determine its leaf area by tracing it on a sheet of paper of known surface density and weighing the leaf’s shape on a fine mass-balance. We filled missing leaf area values with the median value of all leaves of the same species (n = 16).

### Cartridge sampling and analyses

To further identify and quantify the specific monoterpenes emitted from leaves, we collected monoterpenes on glass cartridges filled with Tenax during each pyruvate labeling event. We used an air sampling pump (220-1000TC, SKC, Eighty Four, PA, USA) attached at the outlet to maintain a defined air flow (100 - 200 ml min^-1^) through the cartridge. We attached the cartridges to the cuvette for at least 30 minutes prior to the leaf cutting and for ~70 minutes while the leaf petiole was submersed in the pyruvate solution. Parallel to the collection of each cartridge from the leaf cuvette, we collected a cartridge from the empty cuvette to correct for cuvette background and potential contaminants. After sample collection we sealed the cartridges in vacutainers and stored them at room temperature prior to analysis.

We subsequently analyzed all cartridge samples by gas chromatography – mass spectrometry – combustion interface – isotope ratio mass spectrometry (GC-MS – C – IRMS) at the University of Freiburg to identify and quantify the emitted VOCs and to determine the δ^13^C values of individual VOCs. Volatiles were analyzed after thermodesorption on a GC (7890B, Agilent Technologies Böblingen, Germany) equipped with a mass-selective detector (MSD, 5975C, Agilent Technologies Böblingen, Germany) and a thermodesorption/cold injection system (TDU-CIS4, Gerstel, Germany). VOCs were desorbed from cartridges at 220 °C and immediately trapped in the CIS at −70°C. The CIS was heated to 240 °C, channeling the volatiles onto the separation column (DB-5ms UI, 30 m × 0.25 mm ID, 0.25 μm film thickness, Agilent Technologies Böblingen, Germany). Further details on GC oven and MSD settings are given by Haberstroh et al. (2018). To quantify specific compounds (i.e., main monoterpenes including trans-β-ocimene), we regularly analyzed calibration curves with authentic standards. We analyzed chromatographic raw data with MassHunter Software (Agilent Technologies Böblingen, Germany). We compared mass spectra with the NIST mass spectral library and the retention times of authentic standards to identify compounds of interest.

10 % of the column effluent was diverted to the MSD by a splitter at the end of the GC column, while the remainder of the sample was transferred to an Isotope Ratio Mass Spectrometer (IRMS; Isoprime precisION, Elementar, Hanau, Germany) via a combustion furnace (GC5 interface, Elementar, Hanau, Germany). The latter was operated at 850 °C and combusted VOCs with CuO to CO_2_ and water. After removal of water by a Nafion trap, the CO_2_ was directed into the continuous flow IRMS to analyze compound specific δ^13^C values. We regularly analyzed octane (δ^13^C: −31.75 ± 0.01 ‰) and octadecane (δ^13^C: −32.70 ± 0.01 ‰) (available from Schimmelmann lab, Indiana University, https://arndt.schimmelmann.us/compounds.html) as isotopic reference standards throughout the measurement window. We analyzed raw data with the IonIOS software (Elementar Analysensysteme GmbH, Langenselbold, Germany).

### Calculations

For all calculations, we used 15-minute averages from the end of the labeling period (minute 55-70 after label application) to minimize the effects of any disturbance from cutting the leaf and placing it into solution. We calculated the transpiration rate *E* [mmol m^-2^s^-1^] and the assimilation rate *A* [μmol m^-2^s^-1^] from the concentration of water vapor and CO_2_ as in von Caemmerer and Farquahar (1981), using the concentrations from the stable isotope spectrometers (Picarro L2120i and Thermo Delta Ray, respectively). We calculated instantaneous water use efficiency *WUEi* [μmol CO_2_/mmol H_2_O] as the ratio *A/E*.

We calculated δ^13^C values of CO_2_ relative to the Vienna Pee Dee Belemnite (VPDB) scale using two internal reference gases (δ^13^C values = −9.86 ‰ and −27.8 ‰). We calculated the ^13^C enrichment of CO_2_ (*ε^13^C_CO2_*) leaving the leaf cuvettes following labeling with ^13^C- pyruvate as

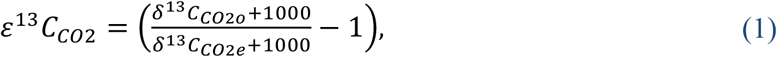

where *δ^13^C_CO2o_* and *δ^13^C_CO2e_* are the δ^13^C values of CO_2_ from the outlet and the entrance of the cuvette, respectively. The term *ε^13^C_CO2_* is functionally equivalent to *Δ^13^C_CO2_* as it is frequently used in plant sciences.

We converted δ^13^C values to the fraction of ^13^C (^13^F = ^13^C/(^12^C + ^13^C)) using the definition of δ^13^C and the known ^12^C/^13^C ratio of VPDB (Hayes, 2004). We then used isotopic mass balance to determine the net isotopic effect of photosynthesis and respiration on the ^13^F value of CO_2_ leaving the leaf cuvette, *^13^F_P_*, prior to labeling as:

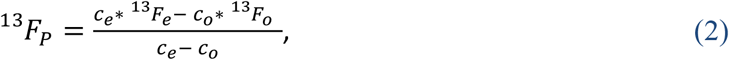

where *^13^F_e_* and *^13^F_o_* are the ^13^F values for CO_2_ and *c_e_* and *c_o_* are the concentrations of CO_2_ at the entrance and outlet of the leaf cuvette, respectively. We averaged *^13^F_P_* values for all measurements of each species, and assumed that it remained constant throughout the pyruvate labeling event, because the natural isotopic effect of photosynthesis and respiration on the δ^13^C value of the CO_2_ leaving the cuvette was relatively small (always less than 10 ‰) relative to the impact of the ^13^C pyruvate label.

We were then able to calculate the concentration of ^13^C originating from the pyruvate label, *c_l_*, through isotopic mass balance as:

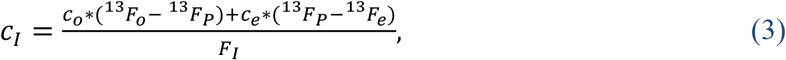

where the ^13^F value of the 99 atom % ^13^C-pyruvate label, *F_l_*, is 0.99. We converted *c_l_* to moles of CO_2_ using the ideal gas law and the flow rate of air to the leaf cuvettes, and to grams of CO_2_ using the molar mass of CO_2_. We then compared this mass to the total mass of ^13^C taken up from the pyruvate label to determine the percent of pyruvate carbon that was allocated to CO_2_.

Volume mixing ratios (VMR, [ppbv]) of VOCs were calculated from counts per second (cps) via mass-dependent transmission of the PTR-TOF-MS, followed by mass scale calibration using PTRwid software (version 003 08-11-2020, (Holzinger, 2015)) following the procedure described by (Holzinger et al., 2019). VOC fluxes, *V_l_*, in [*nmol* m^-2^s^-1^] were calculated from molar flow *u* [*nmol* s^-1^], the leaf area *s* [m^2^], the mixing ratio of the VOC from the outlet of the empty cuvette *v_e_*[*ppbv*], and the mixing ratio of the VOC at the outlet of the leaf cuvette *v_o_*[*ppbv*] as:

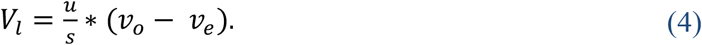

To distinguish VOCs that were synthesized directly from introduced ^13^C1- or ^13^C2- pyruvate, *^13^V_l_*, and VOCs produced only with ^12^C, ^12^*V_l_*, we calculated the fraction of the fluxes of mass 69.0683 in relation to mass 70.0719 for isoprene and mass 137.141 in relation to mass 138.145 for monoterpenes. We subsequently subtracted the theoretical value of the contribution of the heavier isotope under natural abundance of ^13^C (1.1%), ^13^*F_na_* (isoprene: 5.5%; monoterpenes: 11%) to calculate the excess ^13^C fraction ^13^*F_Vl_* in percent:

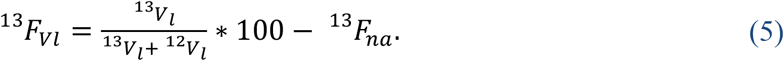

To calculate ^13^C enrichment of individual monoterpenes measured by GC-C-IRMS, we calculated ε^13^C values between the δ^13^C value of the compound during a specific labeling event (δ^13^C_l_) and the mean δ^13^C value of the same compound from unlabeled leaves (δ^13^C_n_) during the same experimental phase (pre-drought or drought):

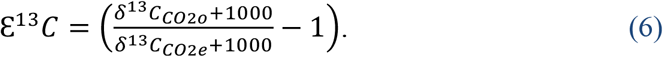

### Statistics

Changes in transpiration, assimilation, VOC fluxes, and ^13^C-enrichment of CO_2_ between pre-drought and drought measurements were compared for each species using Welch’s t test. Differences in ^13^C-enrichment of CO_2_ were also assessed using three-way ANOVA followed by Tukey’s post hoc test, with species, drought treatment, and position of ^13^C in the pyruvate label as separate factors. All statistical analyses were performed in Prism (version 9.5.1, GraphPad Software, LLC).

## Results

### Changes in transpiration, assimilation, and leaf water use efficiency

Transpiration and assimilation rates declined during the drought for all species besides *H. rosa sinensis*, which had low rates in both periods (**Figure 3**). *P. dioica* displayed the highest intraspecies variability in these fluxes, with very high values observed for the leaves of the largest individual, located on the south side of the biosphere, which received more direct sunlight than the individuals on the north side. *C. fairchildiana* displayed the largest relative reductions in transpiration and assimilation, with a relatively greater decline in assimilation, resulting in a significant decrease in water use efficiency (WUE) during the drought (**Figure 3**). Nevertheless, *C. fairchildiana* had the highest WUE of any species in both phases of the experiment (**Figure 3**).

**Figure 3:**
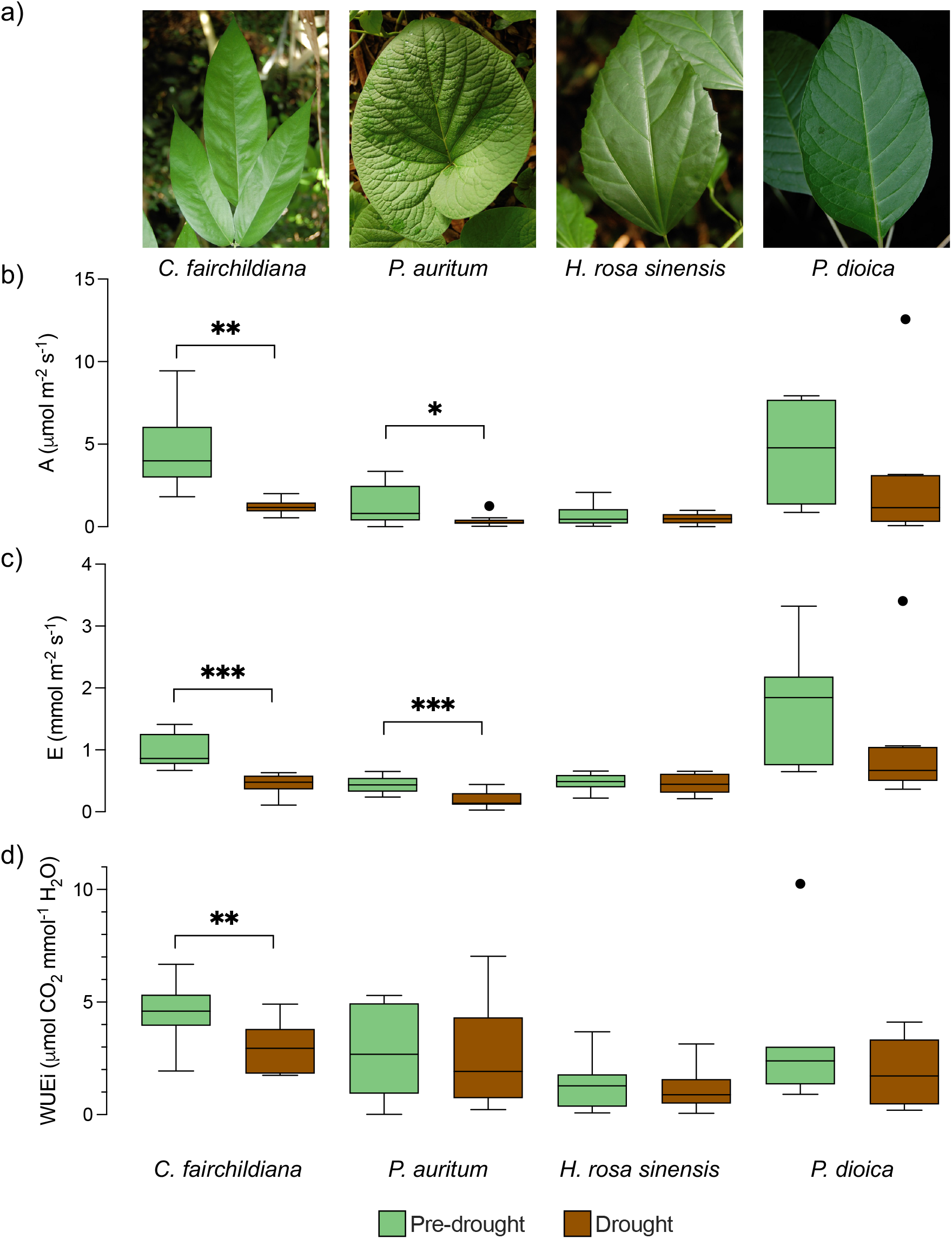
Representative leaves (panel a) and rates of (b) carbon assimilation A, (c) transpiration E, and (d) instantaneous water use efficiency WUEi measured on individual leaves of *Clitoria fairchildiana, Piper auritum, Hibiscus rosa sinensis*, and *Phytolacca dioica* under pre-drought (green) and drought (brown) conditions. Boxes represent the median and 25 – 75 % range of 7-12 replicate leaves. Whiskers and outliers calculated using Tukey’s method. Significant differences between pre-drought and drought conditions for each species are indicated as * when p < 0.05, ** when p < 0.01, and *** when p < 0.001.

### VOC fluxes

*C. fairchildiana* had the highest isoprene fluxes of all species under both pre-drought (7 ± 3 nmol m^-2^ s^-1^) and drought (1.1 ± 0.9 nmol m^-2^ s^-1^) conditions (**Figure 4**). *P. auritum* had the second highest isoprene fluxes, but these were an order of magnitude lower than those from *C. fairchildiana* (0.1 ± 0.04 nmol m^-2^ s^-1^ under pre-drought conditions and 1*10^-4^ ± 9*10^-5^ nmol m^-2^ s^-1^ under drought conditions; **Figure 4**). Isoprene emissions from *H. rosa sinensis* and *P. dioica* were only present in trace amounts (not shown). Between the pre-drought and drought, isoprene emissions only declined significantly for *P. auritum* (p < 0.05), but there were declining tendencies in emissions for *C. fairchildiana* as well (**Figure 4**).

**Figure 4:**
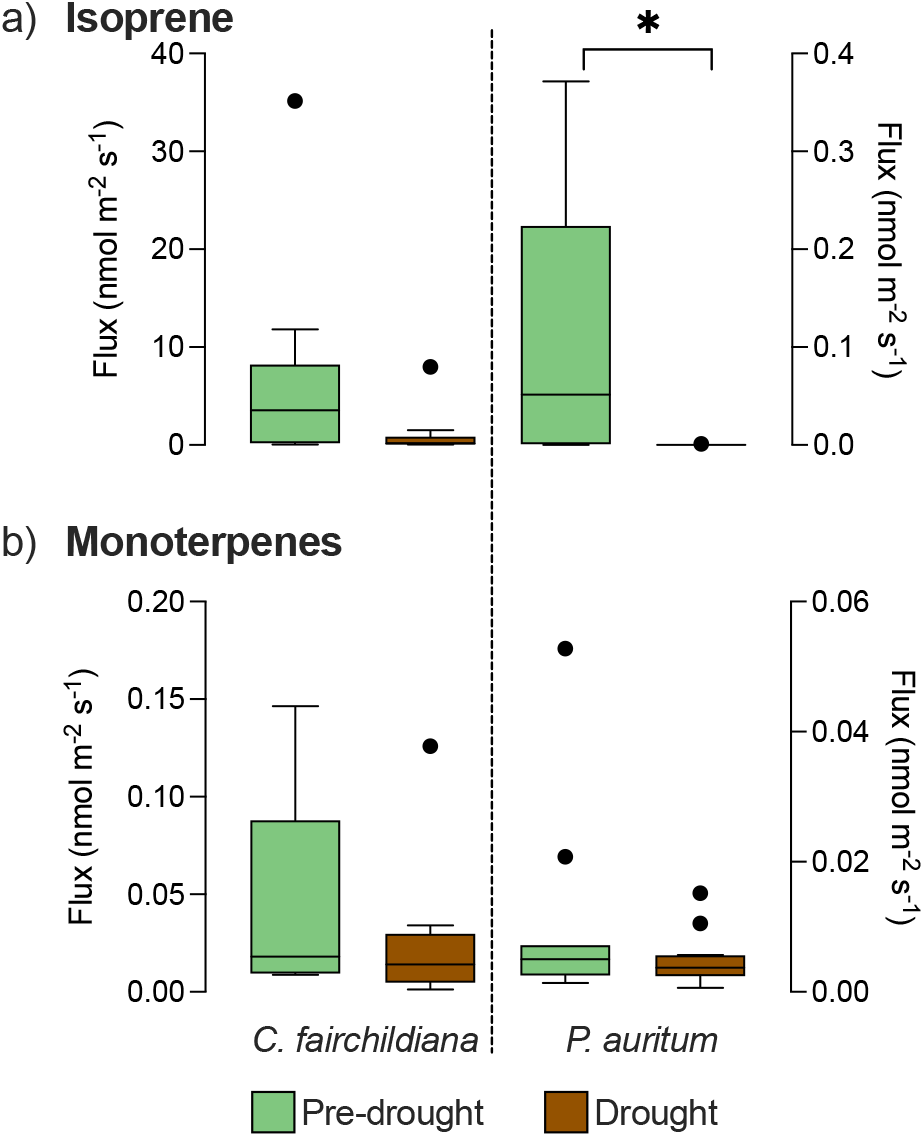
Emission rates of (a) isoprene and (b) monoterpenes measured on individual leaves of *Clitoria fairchildiana* and *Piper auritum* under pre-drought (green) and drought (brown) conditions. Fluxes from *C. fairchildiana* are plotted on the left y-axis and fluxes from *P. auritum* on the right y-axis. Boxes represent the median and 25 – 75 % range of 7-12 replicate leaves. Whiskers and outliers are calculated using Tukey’s method. Significant differences between pre-drought and drought conditions for each species are indicated as: * when p < 0.05.

Monoterpene emissions were also at least an order of magnitude higher from *C. fairchildiana* than any other species (**Figure 4**), but were two orders of magnitude lower than its isoprene emissions. Monoterpene emissions declined under drought (**Figure 4**), but not significantly. Analysis of cartridge samples by GC-MS (which is able to chromatographically resolve different monoterpenes) indicated that a-pinene, β-pinene, 3-carene, limonene, and trans-β-ocimene were the most abundant monoterpenes emitted by *C. fairchildiana* (**Table 1**). Fluxes of most of these compounds declined with drought, but trans-β-ocimene emissions increased by a factor of four (**Table 1**).

**Table 1:**
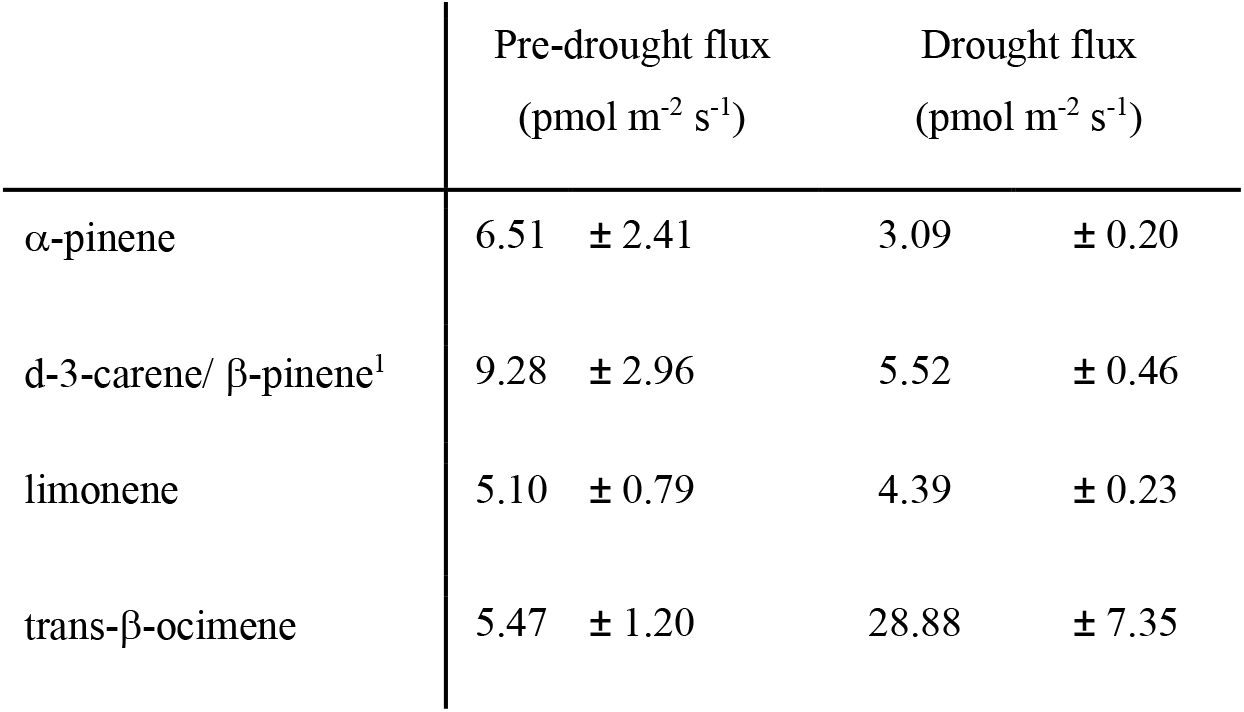
Fluxes of the main monoterpenes emitted by *Clitoria fairchildiana*, as determined by GC-MS analysis. Monoterpene fluxes from the other three species were too low for quantification by GC-MS. Fluxes are reported as mean and standard error. ^1^These two compounds co-eluted on the GC and the combined flux is reported.

|  | Pre-drought flux<br>(pmol m<sup>-2</sup> s<sup>-1</sup>) | Drought flux<br>(pmol m<sup>-2</sup> s<sup>-1</sup>) |
| --- | --- | --- |
| α-pinene | 6.51 ± 2.41 | 3.09 ± 0.20 |
| d-3-carene/ β-pinene<sup>1</sup> | 9.28 ± 2.96 | 5.52 ± 0.46 |
| limonene | 5.10 ± 0.79 | 4.39 ± 0.23 |
| trans-β-ocimene | 5.47 ± 1.20 | 28.88 ± 7.35 |

### ^13^C-enrichment of CO_2_ from pyruvate

All studied plants in Biosphere 2 displayed the expected pattern of more ^13^C-enriched CO_2_ following labeling with ^13^C1-pyruvate than with ^13^C2-pyruvate (**Figures 5, 6**). For most species, the difference in the percentage of ^13^C from ^13^C1-pyruvate that was converted to CO_2_ was significantly higher than the percentage of ^13^C from ^13^C2-pyruvate that was converted to CO_2_ during pre-drought (**Figure 5**). However, this difference was not significant for *C. fairchildiana* at any time, nor for *P. auritum* during drought. As much as ~10 % of the total ^13^C from the ^13^C1-pyruvate was decarboxlated (or released) in the light during the 80 minutes of our measurements, in contrast to less than 1.5 % of the total ^13^C taken up from the ^13^C2- pyruvate (**Figure 5**). However, the magnitude of this difference tended to be less pronounced for *C. fairchildiana* and *P. auritum* than for *H. rosa sinensis* and *P. dioica* (**Figures 5, 6)**.

**Figure 5.**
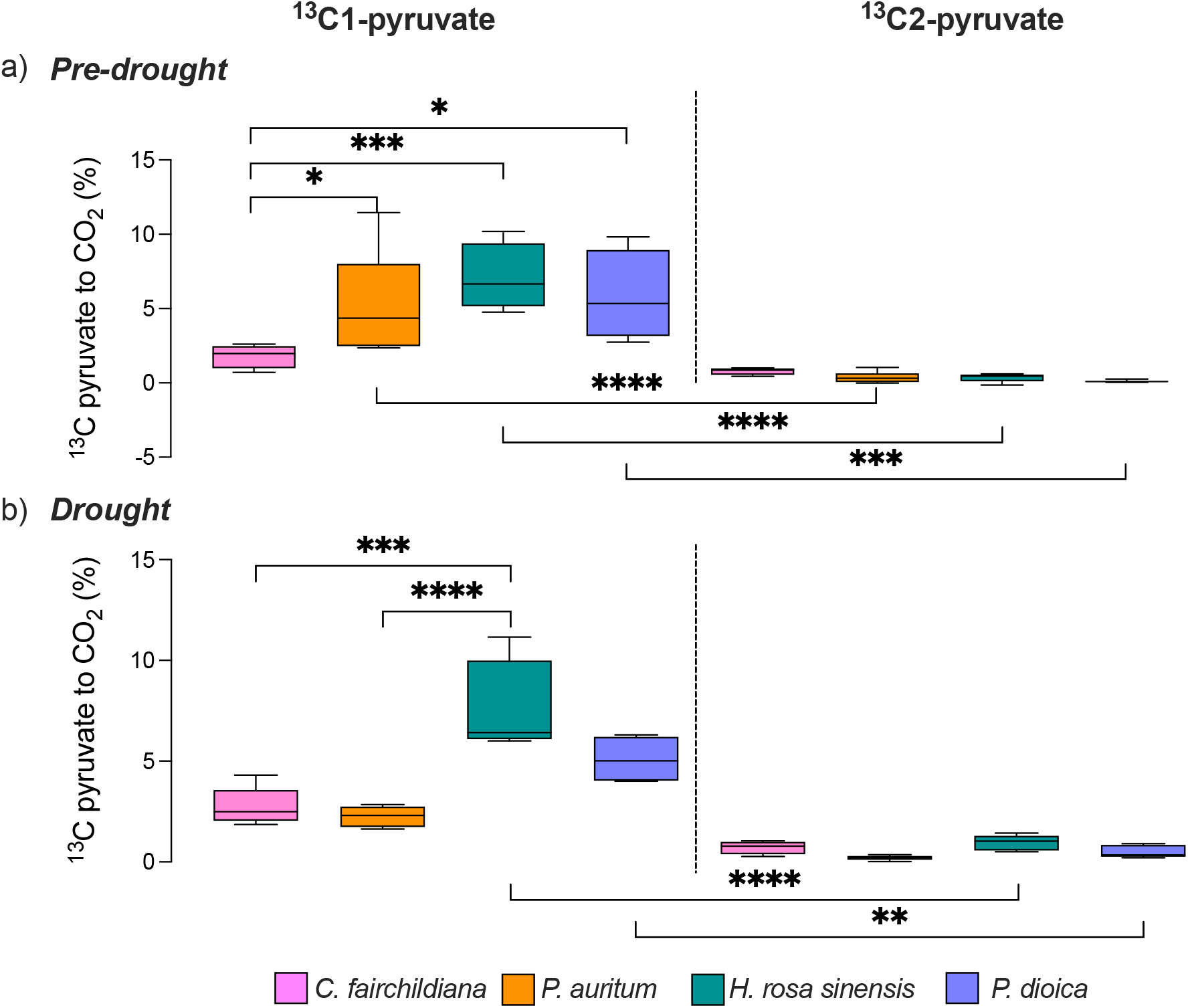
Percent of ^13^C from ^13^C1-labeled pyruvate (left column) and ^13^C2-labeled pyruvate (right column) that was released as CO_2_ by *Clitoria fairchildiana, Piper auritum, Hibiscus rosa sinensis*, and *Phytolacca dioica* under pre-drought (panel a) and drought (panel b) conditions. Boxes represent the median and 25 – 75 % range of 3 - 6 replicate leaves. Whiskers and outliers are calculated using Tukey’s method. Significant differences among groups as determined by three-way ANOVA with Tukey’s posthoc test are indicated as: * when p < 0.05, ** when p < 0.01, *** when p < 0.001, and **** when p < 0.0001.

**Figure 6.**
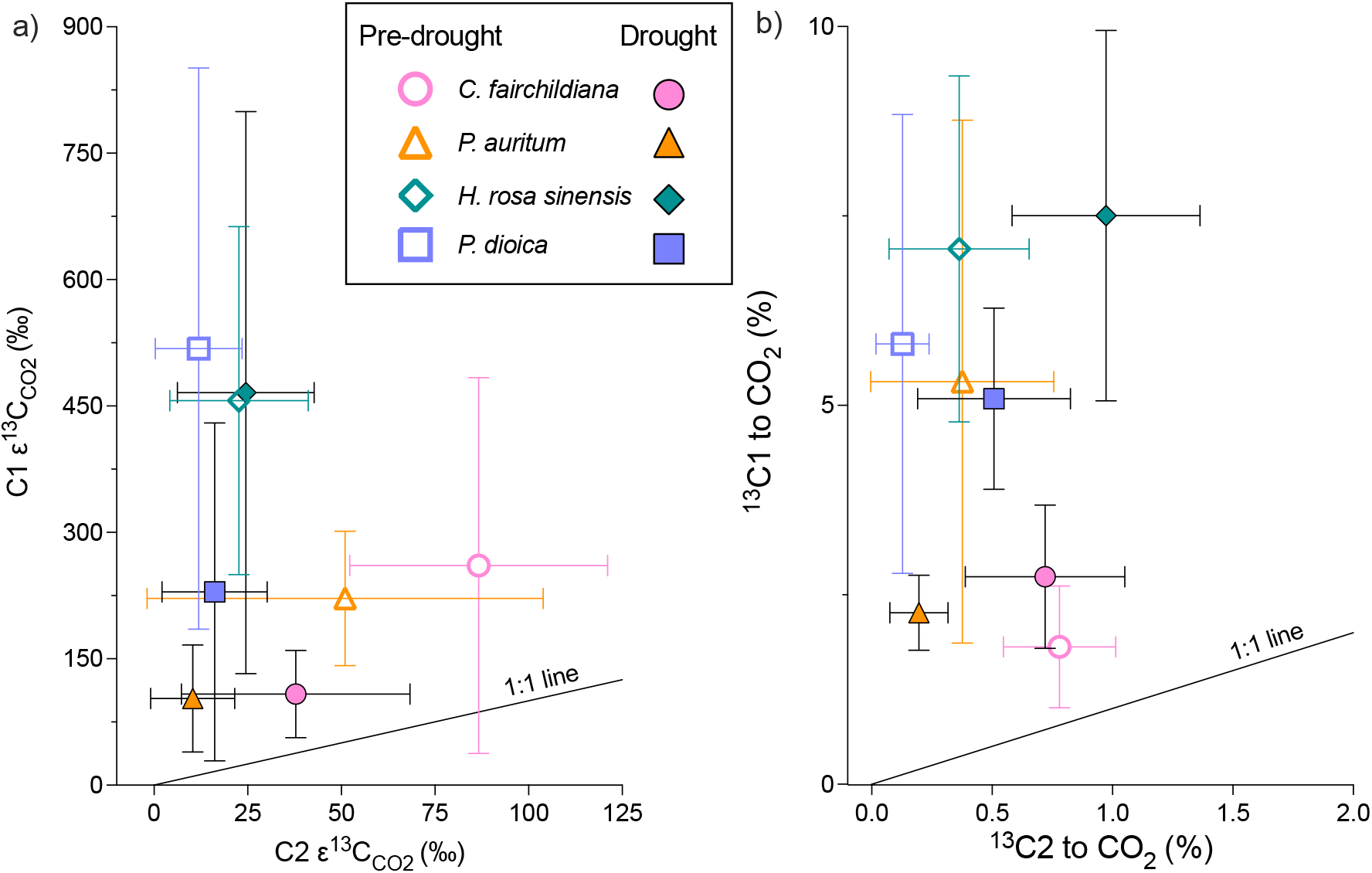
(a) ^13^C enrichment of CO_2_ from ^13^C1-labeled pyruvate (y-axis) plotted relative to ^13^C enrichment of CO_2_ from ^13^C2-labeled pyruvate (x-axis) during pre-drought (open symbols) and drought (closed symbols). (b) Same as panel a, but with the % of ^13^C from each position of the pyruvate label. Symbols represent mean values and error bars represent one standard deviation of 3 – 6 replicate leaves per species.

This difference is apparent by plotting the *ε^3^C_CO2_* (i.e., the change in δ^13^C values of CO_2_ due to labeling) values from ^13^C1-pyruvate labeling relative to those from ^13^C2-pyruvate labeling, where the values for *C. fairchildiana* and *P. auritum* plot much closer to the 1:1 line than the other species do (**Figure 6**). During drought, the *ε^13^C_CO2_* values during labeling with both pyruvate labels tended to decline for both *C. fairchildiana* and *P. auritum*, while for *P. dioica* they only tended to decline for labeling with ^13^C1-pyruvate and for *H. rosa sinensis* they did not change (**Figure 6**).

### ^13^C-enrichment of VOCs following pyruvate labeling

Emissions of isoprene and monoterpenes in *C. fairchildiana* were clearly enriched in ^13^C following the ^13^C-pyruvate labeling during both pre-drought and drought conditions. For isoprene, up to 15 % of the isoprene flux was labeled with ^13^C above the natural background level (**Figure 7a**), indicating that a sizable amount of isoprene was freshly synthesized from pyruvate. During pre-drought conditions, there was no significant difference in the incorporation of ^13^C into isoprene fluxes from the ^13^C1- and ^13^C2-pyruvate labels (**Figure 7a**). Similarly, under drought, there was no significant difference in ^13^C-incorporation between both labels. However, ^13^C-incorporation in isoprene increased under drought, though this was significant only for ^13^C1-labeling (**Figure 7a**).

**Figure 7:**
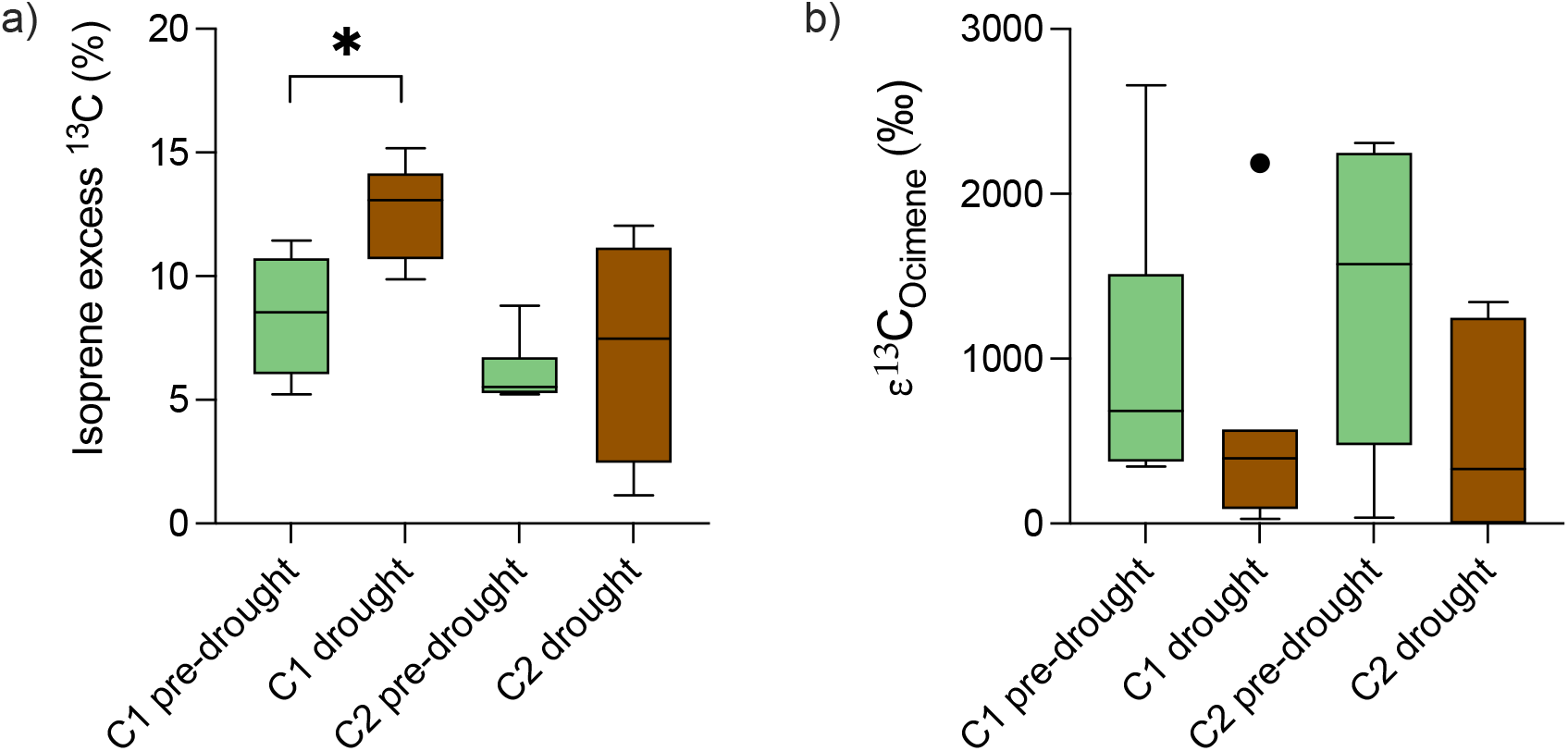
^13^C from ^13^C-labeled pyruvate incorporation into VOCs from *Clitoria fairchildiana*. (a) ^13^C excess isoprene emissions relative to natural background ^13^C abundance for isoprene and (b) ^13^C enrichment of the monoterpene trans-β-ocimene relative to unlabeled measurements during pre-drought (green) and drought (brown) (4 – 6 replicates per treatment). Boxes represent median and 25-75% range of 7-12 replicate leaves. Whiskers and outliers are calculated using Tukey’s method.

Of the monoterpenes emitted from *C. fairchildiana*, only trans-β-ocimene was strongly enriched in ^13^C following labeling by either type of pyruvate, with relatively higher ^13^C enrichment from ^13^C2-pyruvate than from ^13^C1 (**Figure 7b**). Under drought, trans-β- ocimene ^13^C enrichment was similar from both pyruvate labels, with a tendency to decline under drought (**Figure 7b**).

## Discussion

We used position-specific ^13^C-pyruvate labeling to investigate leaf level metabolic processes before and during an experimental drought in the Biosphere 2 tropical rainforest, with an emphasis on understanding changes in allocation of the central metabolite pyruvate into CO_2_ and VOCs in the context of overall drought responses. We focused on four tropical species with clear differences in their drought responses at both the level of individual leaves (**Figure 3**) and in terms of whole plant responses such as leaf water potential, sap flow, and leaf shedding (Werner et al., 2021). In the following discussion, we evaluate our three hypotheses about allocation of ^13^C from pyruvate into CO_2_ and VOCs and place these results into the context of the plants’ overall drought responses.

### Drought response of ^13^CO_2_ production from pyruvate is primarily species dependent

Our first hypothesis, that large reductions in assimilation and transpiration in drought-sensitive plants would also produce less day-time CO_2_ emission from pyruvate, was partially supported. While the drought-tolerant species, *H. rosa sinensis* and *P. dioica*, did not show any decline in the percentage of pyruvate that was converted to CO_2_, the drought-sensitive *P. auritum* did indeed produce less ^13^C-enriched CO_2_ during drought than before (**Figure 5**). Surprisingly, *C. fairchildiana*, the species displaying the overall strongest drought response, did not show such a decline in pyruvate use for ^13^CO_2_ production. Overall pyruvate use differed significantly between *C. fairchildiana* and the other three species (**Figure 5**), indicating fundamental differences in the general use of the central metabolite pyruvate in *C. fairchildiana*, which will be discussed in more detail below.

Our second hypothesis, that the majority of cytosolic pyruvate would be used for non-mitochondrial processes under control conditions, but converge with use in the TCA cycle in accordance to drought response, was also partially supported. In the drought-tolerant species *P. dioica* and *H. rosa sinensis*, cytosolic pyruvate was much more likely to be used for non-mitochondrial anabolic processes than within the TCA cycle before and during drought (**Figure 5, 6**). However, the difference in relative allocation of pyruvate became smaller in *H. rosa sinensis* during drought, mainly because of increased cytosolic pyruvate use within the TCA cycle, as evidenced by increased emissions of ^13^CO_2_ following labeling with ^13^C2- pyruvate (**Figure 5, 6**). This shift towards more daytime TCA activity may have indicated an early drought response in these mostly light-limited plants. On the other hand, for the drought-sensitive *P. auritum* pyruvate consumption declined for all CO_2_ generating processes, indicating an overall slowdown of multiple metabolic pathways with drought. While drought-stressed *P. auritum* plants still tended to produce more ^13^CO_2_ from ^13^C1- pyruvate than from ^13^C2-pyruvate, this difference was no longer significant (**Figure 5, 6**).

Contrary to expectations based on multiple growth-chamber experiments with unstressed plants (e.g., Tcherkez et al., 2008; Priault et al., 2009; Fasbender et al., 2018; Yáñez-Serrano et al., 2019; Werner et al., 2020; Kreuzwieser et al., 2021), the percentage of ^13^CO_2_ that was produced from ^13^C1-pyruvate by *C. fairchildiana* was not significantly higher than that produced from ^13^C2-pyruvate (**Figure 5**). Additionally, the percentage of pyruvate allocated to CO_2_-producing pathways did not change appreciably with drought. Overall, these results suggest that *C. fairchildiana* uses pyruvate in fundamentally different ways than the plants in earlier studies, and that its pyruvate metabolism is minimally affected by drought despite its large drought responses in terms of transpiration and assimilation (**Figure 3**), as well as leaf water potential, sap flow, and leaf shedding (Werner et al., 2021).

### Potential sinks for cytosolic pyruvate in C. fairchildiana

Overall, *C. fairchildiana* decarboxylates significantly lower amounts of cytosolic pyruvate than other plant species, even when it becomes increasingly carbon limited due to strongly reduced assimilation under drought, raising the question how pyruvate is used by this species instead. Here we explore two likely metabolic pathways of use of pyruvate in *C. fairchildiana* that would explain the allocation of all three carbons to biosynthetic products, rather than CO_2_: anaplerotic recapture of decarboxylated C feeding an enhanced, non-cyclic TCA flux that supplies amino acid precursors, and incorporation of the whole pyruvate molecule into isoprenoids via glyceraldehyde 3-phosphate (GA-3-P).

In the light, the reactions of TCA metabolism are better conceived of as a non-cyclic flux rather than a cycle (Sweetlove et al., 2010). While the traditional nocturnal mode of the TCA cycle produces energy and consumes acetyl-CoA produced from pyruvate, this pathway can serve many other functions in plants during the day. In particular, citrate and 2-oxoglutarate produced by TCA metabolism can be exported and form the carbon backbones of amino acids such as glutamate and glutamine (Hanning and Heldt, 1993; Tcherkez et al., 2009). Additional sources of carbon besides acetyl-CoA are required to sustain this carbon flux from the TCA cycle, and can be provided anaplerotically by oxaloacetate (OAA) produced from phospho*enol*pyruvate (PEP) and CO_2_ (Tcherkez et al., 2009; Sweetlove et al., 2013) (**Figure 2**). We hypothesize that the required CO_2_ for OAA production is at least partially directly supplied by the decarboxylation of pyruvate to form acetyl-CoA. This would explain the relatively low amount of ^13^CO_2_ produced from the ^13^C1-pyruvate label that we observed. Indeed, *C. fairchildiana* may have a greater demand for carbon backbones for amino acids than the other species, since it is a legume and therefore has higher nitrogen content than the other taxa. A similar reduction in ^13^CO_2_ production from ^13^C1-pyruvate has also been observed for soil microbial communities when they are provided with succinate or with leaves from legumes, both of which increase nitrogen availability, making an anaplerotically fed open flux mode of the TCA metabolism more favorable (Dijkstra et al., 2011).

Biosynthesis of isoprenoids could be another significant sink for cytosolic pyruvate in *C. fairchildiana*, which was the only strong isoprenoid emitter among the species we analyzed in B2 rainforest (**Figure 4**). Since we observed no differences between ^13^C1- and ^13^C2-pyruvate incorporation into isoprene (**Figure 7**), the primary route by which pyruvate appears to have been incorporated into isoprene in *C. fairchildiana* was via GA-3-P, which conserves all three carbon atoms from the original pyruvate atom, and not via decarboxylated pyruvate (as reported by Yáñez-Serrano et al. 2019; Werner et al. 2020; Kreuzwieser et al., 2021), which would result in greater ^13^C-enrichment following labeling with ^13^C2-pyruvate compared to ^13^C1-pyruvate (**Figure 2**). A significant use of cytosolic pyruvate for isoprenoid biosynthesis via GA-3-P would account for the observed similar production of ^13^CO_2_ from both label types and has previously been observed for some MEP-derived isoprenoids (Ladd et al., 2021). Consequently, these patterns could also partially explain the significant differences in ^13^CO_2_ production from labeled pyruvate that we observed between *C. fairchildiana* and the other three species.

### Different patterns of ^13^C incorporation into isoprene and monoterpenes in C. fairchildiana

We were only able to evaluate our third hypothesis, that the proportion of isoprenoids synthesized from cytosolic pyruvate would increase under drought, for *C. fairchildiana*, which was the only species that emitted sufficiently high enough quantities of isoprenoids for isotopic analyses. In *C. fairchildiana*, this hypothesis was supported for isoprene, as the ^13^C-enrichment of emitted isoprene following ^13^C-pyruvate labeling increased under drought (**Figure 7**), even while overall isoprene emissions declined (**Figure 4**). These results indicate that synthesis of isoprene from primary photosynthate by *C. fairchildiana* declined under drought as assimilation also declined, resulting in an increased proportion of the remaining isoprene flux being dependent on cytosolic pyruvate (**Figure 8, step 2-4**) similar to what has been observed for heat-stressed plants in growth chambers (Yáñez-Serrano et al., 2019).

**Figure 8:**
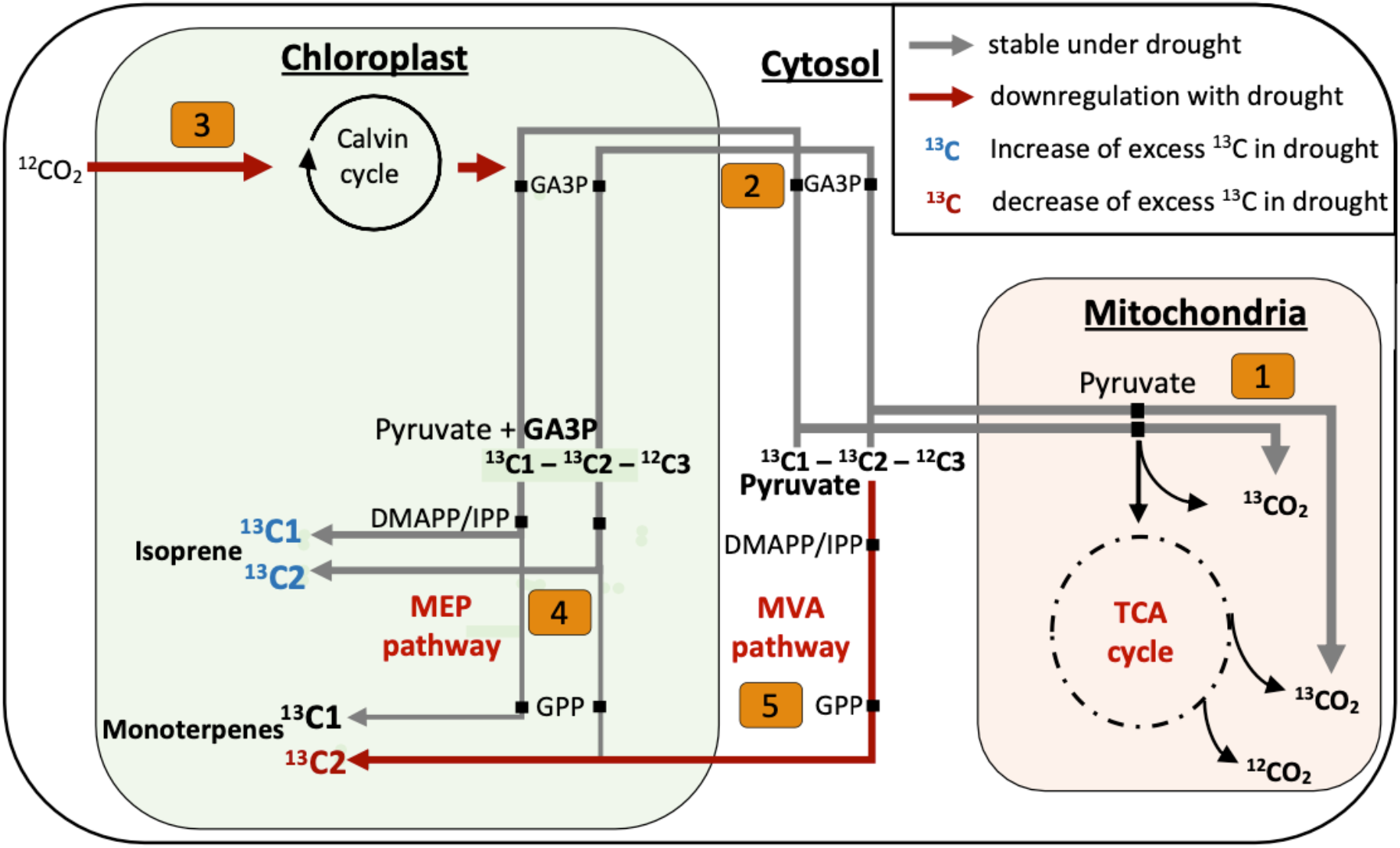
Schematic representation of pyruvate carbon partitioning among biosynthetic processes involved in the synthesis of isoprenoids and production of CO_2_ within *Clitoria fairchildiana* cells as well as proposed adjustments of these processes during drought. Several intermediate steps and involved enzymes are removed for clarity. Arrow thickness represents the relative amount of ^13^C1-/ ^13^C2-labeled pyruvate used for the indicated pathway during pre-drought conditions. Gray arrows indicate no changes in carbon utilization during drought while brown arrows indicate a down-regulation of the pathway. Blue colored ^13^C1/^13^C2 indicate an increase in excess ^13^C emissions of isoprene or monoterpenes during drought, while red color indicates a decline in excess ^13^C emissions. Numbers represent the relevant pathways discussed, with descriptions of pathways 1-5 described in the caption of Figure 1. TCA, tricarboxillic acid; GA-3-P, glyceraldehyde 3-phosphate; DMAPP Dimethylallyl pyrophosphate; IPP, isopentenyl pyrophosphate; MEP, methylerythritol phosphate; GPP, geranyl pyrophosphate; MVA, mevalonic acid.

Surprisingly, increased ^13^C-enrichment of monoterpenes following ^13^C-pyruvate labeling did not increase under drought (**Figure 7**), contradicting our expectations that isoprene and monoterpenes would behave similarly to each other since they are both derived from IPP and its isomer dimethylallyl pyrophosphate (DMAPP) produced in the MEP pathway (**Figure 2, step 4**). IPP and DMAPP are also produced in the cytosolic MVA pathway, where pyruvate is incorporated only via acetyl-CoA, with its C1 position being decarboxylated (**Figure 2, step 5**). Possibly, geranyl pyrophosphate (GPP), an intermediate between IPP and monoterpenes that is produced in both the MVA and MEP pathways (Bouvier et al., 2001; Schmidt and Gershenzon, 2007; Zulak et al., 2010), is produced from cytosolic pyruvate and transported into the plastid to supply monoterpene synthesis, but not isoprene synthesis (Bick and Lange, 2003; Gutensohn et al., 2012) during pre-drought conditions. This would explain why pre-drought trans-β-ocimene tended to become more ^13^C-enriched following labeling with ^13^C2-pyruvate than with ^13^C1-pyruvate while the relative ^13^C-enrichment was similar for isoprene from both labels (**Figure 8**). Alternatively, the same pattern could be observed if trans-β-ocimene was synthesized as a byproduct of sesquiterpene synthesis (Davidovich-Rikanati et al. 2008; Gutensohn et al. 2013) or via enzymes of the Nudix hydrolase family (Magnard et al., 2015; Liu et al., 2018) in the cytosol. However, these processes are unlikely to drive the observed patterns as *C. fairchildiana* did not emit any sesquiterpenes and the presence of monoterpene synthesis via Nudix hydrolases in vegetative tissue has yet to be shown. We therefore consider an MVA-derived GPP contribution to pre-drought monoterpenes to be a more likely explanation for the different patterns of ^13^C incorporation into isoprene and trans-β-ocimene.

During drought, trans-β-ocimene synthesis from MVA derived GPP apparently declined or even completely ceased (**Figure 8, step 5**), as the ^13^C-enrichment of β-ocimene was similar from both ^13^C1- and ^13^C2-pyruvate, similar to isoprene (**Figure 7**). In contrast to isoprene, ^13^C enrichment of trans-β-ocimene tendentially declined with drought, indicating less reliance on cytosolic pyruvate for the synthesis of these compounds, even as cytosolic pyruvate apparently became more important for supporting the isoprene flux (**Figure 7**). The differing patterns for ^13^C incorporation into isoprene and trans-β-ocimene emissions are intriguing and further experiments are needed to fully disentangle the mechanisms responsible for the observed patterns.

## Conclusions

Biochemical experiments including position-specific ^13^C-labeling are usually conducted under highly-controlled laboratory conditions on small saplings. Here, we applied this approach to mature plants in a near-natural ecosystem. Three of the four species we investigated generally conformed to expectations about the relationship between functional drought responses and use of cytosolic pyruvate. Specifically, daytime production of CO_2_ from pyruvate declined more as drought stress increased, more daytime CO_2_ was produced from non-mitochondrial biosynthetic pathways than from TCA metabolism, and there was convergence between these two sources of CO_2_ as drought stress increased. However, we observed strikingly different and novel results from *C. fairchildiana*, a legume tree that is also a strong isoprenoid emitter. In particular, CO_2_ production from pyruvate was relatively low in this species during both pre-drought and drought conditions, and CO_2_ production from the C1 position of pyruvate was not significantly higher than from the C2 position. This suggests that *C. fairchildiana* efficiently incorporates all three carbon atoms from pyruvate into metabolic products including amino acids and isoprenoids. The patterns of ^13^C incorporation into different isoprenoids emitted by *C. fairchildiana* also suggests that crosstalk between the cytosolic MVA and the plastidic MEP pathways for isoprenoid synthesis is important for monoterpene synthesis, but not isoprene synthesis, and that this metabolic crosstalk declines with drought. *C. fairchildiana* over-proportionally contributed to overall isoprene emissions and GPP of the whole ecosystem before drought and played a major role in the decline of these fluxes during drought (Werner et al., 2021). Identifying and understanding metabolic adjustments of drought-sensitive plant species like *C. fairchildiana* that significantly contribute to ecosystem VOC emissions in natural tropical ecosystems could therefore help to improve current attempts to model drought impacts on VOC production in tropical rainforests, which are the major source (>80 %) of global isoprene emissions (Sindelarova et al., 2014).

## Supplementary data

**Table S1:**
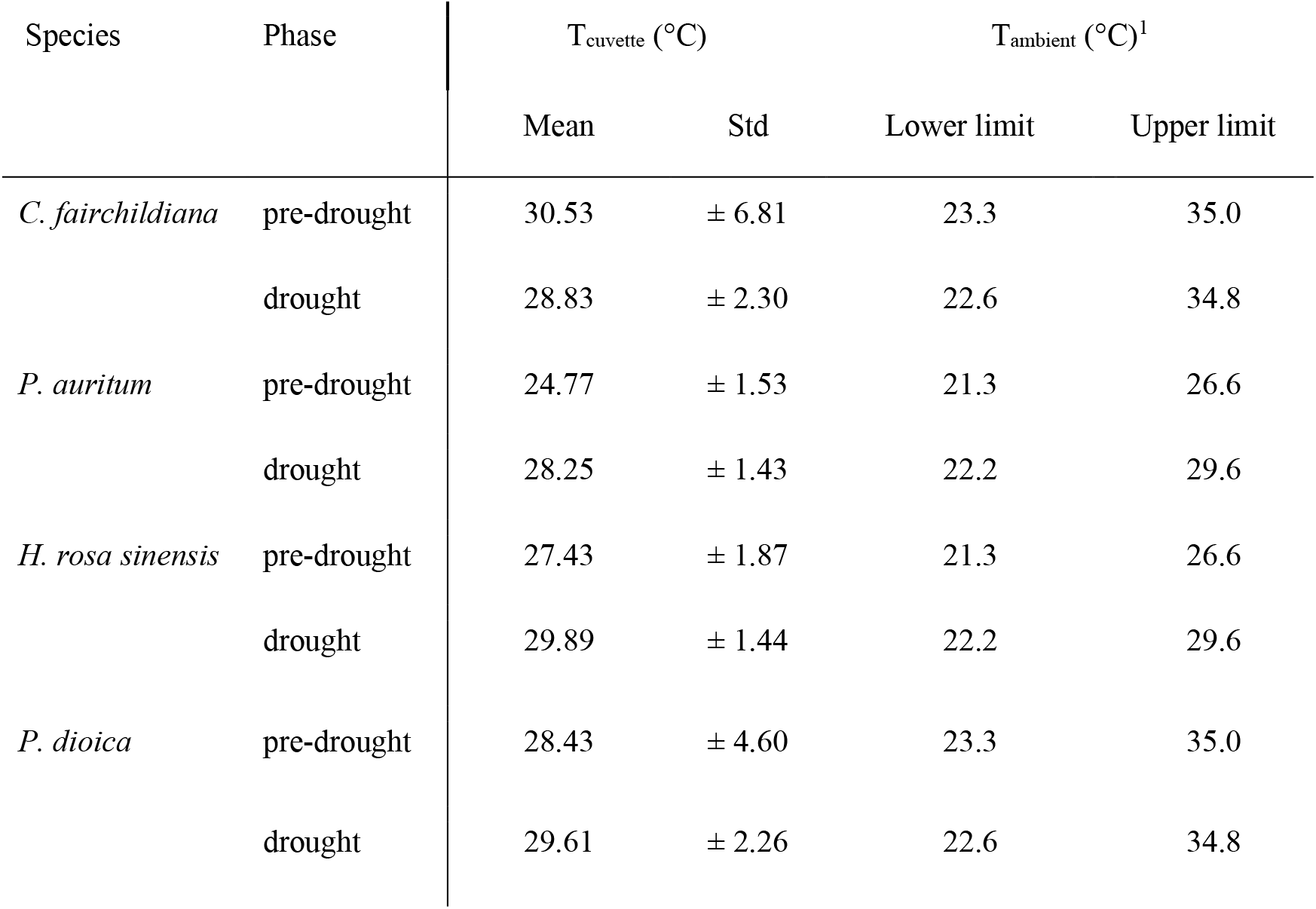
Leaf temperature inside enclosed leaf cuvettes during the experiments in comparison to ambient daytime temperature ranges inside the Biosphere 2 tropical rainforest. ^1^ Data from Werner *et al*., 2021.

## Acknowledgements

The study took place within the context of the B2WALD campaign, and was supported by the wider efforts of the entire B2WALD team, as described on the team contribution list (https://doi.org/10.25422/azu.data.14632662). We are particularly grateful to Michael Burman, Marissa Clover, Sydney Kerman, and Luke Miller for their assistance with leaf labeling in the forest canopy.

## Author Contributions

Conceptualization (CW, SNL, LED); Data curation (LED, IB, JVH, LKM), Formal analysis (LED, SNL, JK, IB, AK, JVH, GP); Funding acquisition (CW, LKM); Investigation (SNL, LED, IB, JD, JVH, GP, JI, CW, AK); Methodology (CW, JK, IB, LED, SNL); Project administration (CW, SNL); Resources (CW, JVH, LKM); Supervision (CW, LKM, SNL); Validation (AK, CW, IB, LED, SNL); Visualization (LED, SNL); Writing – original draft (SNL, LED); Writing – review & editing: all.

