## Supplemental Table 1 for "Leaf-level metabolic changes in response to drought affect daytime CO_2_ emission and isoprenoid synthesis pathways"

**Contents:**

**Table S1**

**Table S1:** Leaf temperatures inside enclosed leaf cuvettes during the experiments in comparison to ambient daytime temperature ranges inside the Biosphere 2 tropical rainforest. [1] Data from Werner *et al.*, 2021.

| Species | Phase | $T_{cuvette}$ (°C) | | $T_{ambient}$ (°C)[1] | |
|---|---|---|---|---|---|
| | | Mean | Std | Lower limit | Upper limit |
| *C. fairchildiana* | pre-drought | 30.53 | ± 6.81 | 23.3 | 35.0 |
| | drought | 28.83 | ± 2.30 | 22.6 | 34.8 |
| *P. auritum* | pre-drought | 24.77 | ± 1.53 | 21.3 | 26.6 |
| | drought | 28.25 | ± 1.43 | 22.2 | 29.6 |
| *H. rosa sinensis* | pre-drought | 27.43 | ± 1.87 | 21.3 | 26.6 |
| | drought | 29.89 | ± 1.44 | 22.2 | 29.6 |
| *P. dioica* | pre-drought | 28.43 | ± 4.60 | 23.3 | 35.0 |
| | drought | 29.61 | ± 2.26 | 22.6 | 34.8 |
